# The biaxial mechanics of thermally denaturing skin - Part II: Modeling

**DOI:** 10.1101/2021.06.04.447120

**Authors:** Manuel Rausch, William D. Meador, John Toaquiza Tubon, Omar Moreno-Flores, Adrian Buganza Tepole

**Affiliations:** Department of Biomedical Engineering, The University of Texas at Austin, Austin, TX, 78712, USA; Department of Aerospace Engineering and Engineering Mechanics, The University of Texas at Austin, Austin, TX, 78712, USA; Oden Institute for Computational Engineering and Sciences, The University of Texas at Austin, Austin, TX, 78712, USA; School of Mechanical Engineering, Purdue University, West Lafayette, IN, 47907, USA; Weldon School of Biomedical Engineering, Purdue University, West Lafayette, IN, 47907, USA

## Abstract

Understanding the response of skin to superphysiological temperatures is critical to the diagnosis and prognosis of thermal injuries, and to the development of temperature-based medical therapeutics. Unfortunately, this understanding has been hindered by our incomplete knowledge about the nonlinear coupling between skin temperature and its mechanics. In Part I of this study we experimentally demonstrated a complex interdependence of time, temperature, direction, and load in skin’s response to superphysiological temperatures. In Part II of our study, we test two different models of skin’s thermo-mechanics to explain our observations. In both models we assume that skin’s response to superphysiological temperatures is governed by the denaturation of its highly collageneous microstructure. Thus, we capture skin’s native mechanics via a microstructurally-motivated strain energy function which includes probability distributions for collagen fiber orientation and waviness. In the first model, we capture skin’s response to superphysiological temperatures as a transition between two states that link the kinetics of collagen fiber denaturation to fiber coiling and to the transformation of each fiber’s constitutive behavior from purely elastic to viscoelastic. In the second model, we capture skin’s response to super-physiological temperatures instead via three states in which a sequence of two reactions link the kinetics of collagen fiber denaturation to fiber coiling, followed by a state of fiber damage. Given the success of both models in qualitatively capturing our observations, we expect that our work will provide guidance for future experiments that could probe each model’s assumptions toward a better understanding of skin’s coupled thermo-mechanics and that our work will be used to guide the engineering design of heat treatment therapies.

## Introduction

Life is highly temperature sensitive with even the most well-adapted metazoans succumbing to temperatures beyond 50 °C [1]. Indeed, mammals thermo-regulate within a very narrow body temperature range of 35-40C. Beyond these temperatures, vital proteins, such as the ubiquitous structural protein collagen, thermally denature [2]. That is, collagen’s complex hierarchical organization disassociates. As collagen breaks down, so do the tissues whose microstructure is governed by collagen [3, 4]. The exact mechanisms that link collagen denaturation to those tissues’ thermo-mechanical response to superphysiological temperatures are not fully understood. However, our current understanding is that the breaking of collagen’s inter- and intra-molecular cross-links from exposure to superphysiological temperatures leads to an energetically more favorable coiled state, i.e. a reduction in the collagen molecule reference length. Loss of cross-links also leads to a reduced load-bearing capacity of collagen [5, 6, 7]. As a result, at the macroscopic scale, superphysiological temperature loading shrinks collageneous tissues and changes their characteristic J-shape stress-strain curve into a more linear behavior. Ultimately, once collagen is fully denatured, collageneous tissues take a gelatinous form and lose their mechanical integrity completely [8, 9, 10].

Collagen is a critical constituent that supports connective tissues such as skin. In turn, understanding thermal denaturation of skin is important to understanding the pathophysiology of thermal injury, such as the one million burn wounds that are treated each year in the US alone [11, 12]. Understanding skin’s response to superphysiological temperatures is also important for the development of temperature-dependent therapeutics. For example, laser therapy has emerged as an effective treatment for scar tissue [13]. Despite the importance of understanding skin’s response to thermal loading, many gaps in knowledge remain. Among them, we currently lack comprehensive knowledge of the relationship between superphysiological temperatures and skin’s mechanical behavior. In Part I of this study, we aimed at filling some of these gaps and concluded that thermal treatment of skin above 55°C results in time-, temperature-, and direction-dependent shrinkage under isotonic traction-free conditions. Similarly, we found that thermal treatment of skin above 55C results in time-, temperature-, and direction-dependent force production under isometric conditions. Additionally, we concluded that thermal treatment of skin results in stiffening and softening of skin at low and high strains, respectively. Supported by histology and second harmonic generation imaging, we also demonstrated that these thermo-mechanical phenomena are driven by the denaturation of collagen. Our work thus confirmed and built on others’ previous work on the thermo-mechanical coupling of denaturing, collageneous soft tissues. For example, Chen at al. subjected chordae tendineae to different thermo-mechanical loading conditions and observed that tissue shrinkage and force production demonstrated time-temperature-load equivalence [10, 14]. Similarly, Harris et al. and Baek et al. found that epicardium exhibited a less pronounced strain-stiffening response after thermal treatment than before [15, 9]. Similar observations were made by Boronyak et al. and Bender et al after radio-frequency ablation of valvular tissue [16, 17]. Finally, skin-specific work by Zhou et al. and Xu et al. showed, comparably to our findings, that the modulus of the tissue decreases after thermal treatment and that its viscoelastic response became more pronounced [18, 8]. Capturing above observations in quantitative models is a critical element in generating and testing hypotheses that can explain the origins of these observations.

There have been previous efforts toward such quantitative models [19, 18, 20]. While each of those models captured some elements of the complex, coupled interplay between thermal treatment and mechanics, none can comprehensively explain time-, temperature, direction-, and load-dependent observations that experimental studies have identified. Part II of our study tests competing models of skin’s response to superphysiological temperatures that can explain ours and others’ recent experimental observations on the coupled thermo-mechanics of denaturing collageneous tissues with a specific focus on skin [21]. Our models build on the the well-established Arrhenius-type kinetics of collagen denaturation [22]. We specifically aim at linking these kinetics to denaturation-induced changes on the tissue scale such as shrinkage, change in mechanical properties, change in viscoelastic properties and/or damage. To this end, we extend previous modeling efforts and propose two competing models. In both models, we consider collagen the primary, structural constituent of skin. Thus, we assume that it is the denaturation of collagen that drives the thermo-mechanical response of skin to superphysiolgocal temperatures. First, we introduce a constitutive model of intact skin that captures tissue’s nonlinear and anisotropic response by explicitly modeling the collagen fiber orientation and waviness distributions. In our first model, we consider collagen, and thus skin, denaturation as a two-state phenomenon where the denatured collagen changes its reference configuration and becomes viscoelastic. In our second model, we consider collagen denaturation as a three-state phenomenon in which no viscoelasticity is assumed. Instead, the denaturation process is split into two steps: an intermediate state of collagen molecules with a reduced stress-free length before a final, damaged state in which the collagen molecules cannot carry mechanical load. Both implementations are schematized in Figure 1.

**Figure 1:**
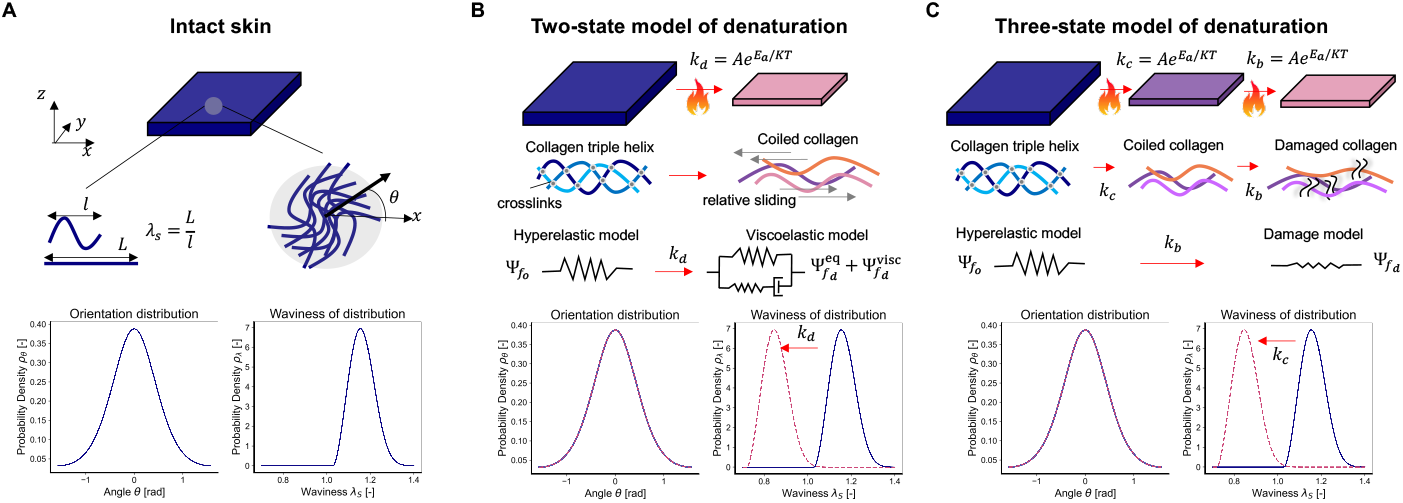
Competing models of skin’s thermo-mechanical response to superphysiological temperatures. A) Independent of the model, we approximate intact skin’s constitutive behavior as hyperelastic, where the strain energy function is microstructurally-inspired by explicitly considering probability distributions of collagen orientation and waviness. B) The two-state model predicts thermally-induced changes in skin’s mechanical properties via one reaction that simultaneously induces two changes: i) a transition in the stress-free reference configuration toward a smaller (coiled) collagen length which is captured by a translation of the waviness distribution, ii) inclusion of viscoelasticity in the collagen behavior which we argue is due to inter- and intra-fibrillar cross-links disassociating, thus, enabling relative sliding within and between fibers. C) The three-state model predicts thermally-induced changes in the mechanical properties of denaturing skin via two subsequent reactions: i) initially, we argue that denaturation leads to coiling of collagen molecules that we, again, capture through a translation of the collagen waviness distribution, ii) in a second reaction, we argue that continued disassociation of cross-links leads to loss of collagen’s strain energy storage capacity, i.e., reduction in material stiffness or damage.

## Materials and Methods

### Mechanics of intact skin

We model the native state of skin as a hyperelastic, constrained mixture with contributions to the strain energy from an isotropic matrix, *W* ^m^, and distributed collagen fibers, 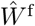. To this end, we characterize the local deformation by the deformation gradient **F** and express the strain energy as a function of the right Cauchy-Green deformation tensor **C**= **F**^⊤^**F**. Specifically, we capture the contribution of the isotropic matrix via the deformation tensor’s first invariant *I*_1_ = **C**: **I**. In contrast to the common approach of capturing collagen’s contribution to the strain energy via a condensation into a single pseudo-invariant [23, 24], we choose a microstructurally-inspired approach [25, 26, 27]. The contribution of a single fiber in the **a**_0_’s material direction to the strain energy is through its pseudo-invariant *I*_4_(*θ*) = **C**: **a**_0_(*θ*). The total contribution of all fibers to the strain energy, 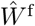, thus follows from integration across the contributions of each individual fiber at angle *θ*, *W* ^f^ (*θ*), according to their probability density distribution *ρ^θ^*(*θ*). Throughout this work, we also assume that skin behaves incompressibly, which we enforce via the Lagrange-multiplier *p*. The total strain energy function consequently reads

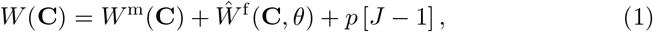

with

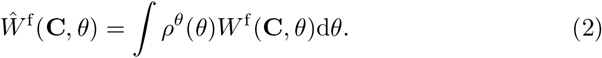

Here we approximate the matrix behavior as neo-Hookean

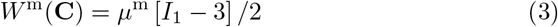

with *μ*^m^ being its shear modulus. In addition to capturing collagen’s distributed orientation, we also capture the distribution of its waviness, which is a microstructural characteristic of collagen in skin and other soft tissues. To this end, we introduce the waviness distribution *ρ^λs^*(*λ_s_*), where *λ_s_* may be interpreted as a distributed reference length [28, 27]. That is, while the fiber is crimped we assume it does not provide resistance to stretch and only does so once uncrimped. Consequently, the stretch-induced energy contribution of each fiber, *W* ^f^, follows from its relative orientation, via *θ*, and its individual reference length, via *λ_s_*, viz.

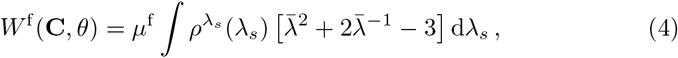

where *λ*=*λ*(**C***, θ*) is the macroscopic stretch in the *θ*-direction, i.e., the square root of the pseudo-invariant *I*_4_(*θ*) = **C**: **a**_0_(*θ*), and 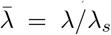 is the fiber stretch relative to its respective reference length. The description of each fiber’s strain energy contribution is that of a neo-Hookean cylinder under uniaxial extension and with traction-free boundaries. In our model, each collagen fiber’s constitutive behavior is thus determined by a single material parameter, its shear modulus *μ*^f^. However, its contribution to the total strain energy depends also on its orientation and its reference length.

The second Piola-Kirchhoff stress tensor for intact skin then follows from the Clausius-Duhem inequality as

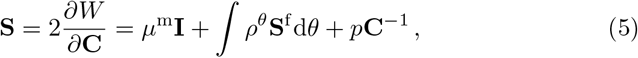

where we dropped the dependencies on *θ* and **C** for convenience. The fiber contribution to the stress for each fiber, in turn, reads

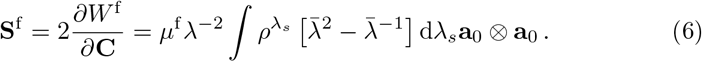

Please note that we model skin as a thin membrane that fulfills the plane-stress condition. Accordingly, we model fibers only within this plane. Additionally, this assumption implies zero normal surface tractions and allows us to analytically solve for the Lagrange multiplier *p*.

For the orientation distribution function, *ρ^θ^*, we consider a *π*-periodic von Mises distribution

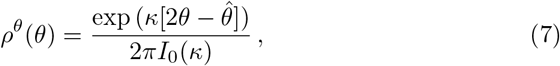

with concentration parameter *κ*, the mean fiber orientation 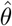, and with *I*_0_(*κ*) being the Bessel function of order zero. For the waviness distribution function, 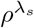, we use the Weibull distribution

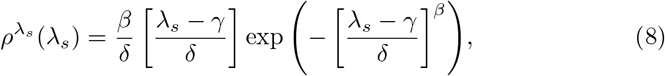

where the parameter *γ* controls the shift of the distribution along the abscissa, while *β* and *δ* are called the shape and scale parameters and control the skewness and width of the distribution, respectively. The orientation and waviness distribution for intact skin are illustrated in Figure 1A.

As a first approximation for above parameters, we informed *μ*^m^ and *μ*^f^ from our previous work [29] and manually adjusted the distribution function parameters to reflect our baseline experiments on dorsal murine skin in Part I, see Table 1. Note that we chose 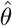 of the von Mises distribution equal to zero (i.e., lateral direction collagen).

**Table 1:** Constitutive parameters of intact skin

| Matrix<br>Stiffness | Collagen<br>Stiffness | Orientation<br>Parameters | Waviness<br>Parameters |  |  |
| --- | --- | --- | --- | --- | --- |
| $\mu^m$ [MPa] | $\mu^f$ [MPa] | $\kappa$ [-] | $\beta$ [-] | $\delta$ [-] | $\gamma$ [-] |
| 0.001 | 1.25 | 2.6 | 0.15 | 1.03 | 0.5 |

### Mathematical models of skin’s response to superphysiological temperatures

In the design of both modeling approaches we set out to capture three heat-induced phenomena that we observed in Part I of this study. When heated, i) skin shrinks under traction-free conditions, ii) skin produces tractions that quickly decay when under isometric conditions, and iii) skin’s constitutive behavior becomes stiffer at low strains and softer at high strains. The below models are built on competing assumptions as to the molecular mechanisms that lend them the ability to predict these phenomena. However, both models share the assumption that collagen and, therefore, collageneous soft tissue denature according to first order kinetics with Arrhenius-type rate, viz.

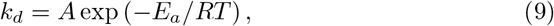

where *k_d_* is the rate of denaturation, *A* is the maximum rate, *E_a_* is the activation energy, *R* is the universal gas constant, and *T* is the temperature. This assumption is based on previous work by Xu et al., for example, who measured *E_a_* = 5.867 × 10^5^J/mol and *A* = 5.240 × 10^91^s^−1^ [8], while Stylianopoulos identified *A* = 1.1363 × 10^86^s^−1^ and *E_a_* = 5.6233 × 10^5^J/mol [20], and Chen et al. measured *A* = 1.3 10^53^s^−1^ and *E_a_* = 3.57kJ/mol; see also the excellent review by Wright et al. [22].

### Two-state model

In our first model, we propose that denaturation of collagen is governed by a transition from a native state to one denatured state. This second, non-native state is characterized by a reduction in collagen’s reference (coiled) length and a transition from being a purely elastic material to being a viscoelastic material, see Figure 1B. The first change in collagen’s properties is motivated by the breaking of inter- and intra-molecular crosslinks that leads to the dissociation of the collagen triple helix and coiling of collagen molecules; we capture this change by a change in the fibers’ waviness distribution. The second change in collagen’s properties is motivated by the breaking of inter- and intra-molecular crosslinks that leads to relative sliding between collagen fibrils under load; we capture this change by altering the fibers’ hyperelastic constitutive model to a hyper-viscoelastic one. In turn, the transition between both states is governed by first order kinetics, see above. That is, change of the concentration of native collagen, *C_o_*, to denatured collagen, *C_d_*, is governed by the rate *k_d_*. The strain energy of the mixture thus reads

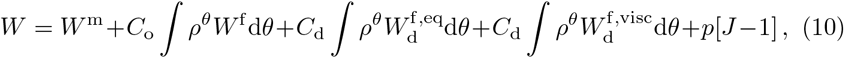

where the subscript ‘d’ denotes quantities associated with denatured collagen and tissue. For example, 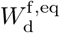 and 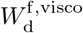 represent the damaged fibers’ equilibrium contribution and a viscoelastic contributions to the strain energy function. Specifically,

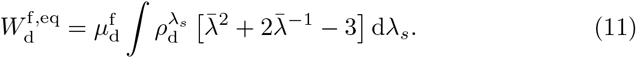

Here, again, note that the subscript ‘d’ in 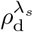 and 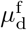 indicates that those quantities change during denaturation (4). Analogous to the equilibrium contribution, the viscoelastic contribution reads

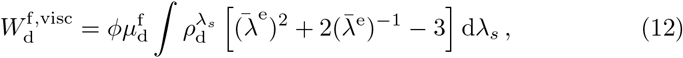

except that its evolution is driven by the elastic deformation of a Maxwell-type branch 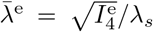 rather than the total stretch *λ*. Note, following [30, 31] it is assumed that the viscoelastic part of the strain energy function is proportional to the equilibrium branch through the factor *ϕ*. The second Piola Kirchhoff stress tensor then follows from the Doyle-Erickson relation as

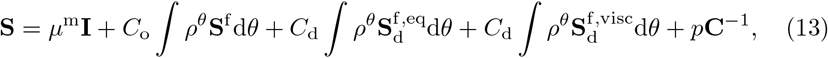

where the fiber equilibrium and viscoelastic stresses, 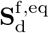 and 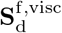, respectively, are computed as in Equation 6. For the update of the viscous strain, consider that the rate of change of the stress is proportional to the stress,

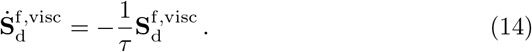

As a result, the rate of change of the viscous strain follows from solving

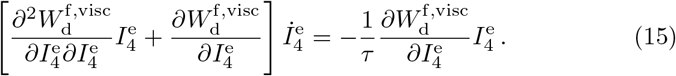

For more details on this hyper-viscoelastic formulation we refer the reader to Liu et al. [30]. Additionally, the parameters for the two-state model are summarized in Table 2. Note that the parameters for the matrix contribution and the orientation distribution remain unchanged from the intact tissue properties, see Table 1. Furthermore, the parameters in Table 2 are estimated based on previous work on modeling collagen denaturation, e.g. [10, 14, 20], and manual adjustment to reproduce qualitatively the observations from Part I [21].

**Table 2:** Parameters for the two-state model of skin’s thermo-mechanical response to super-physiological temperatures

| Kinetics |  | Waviness |  |  | Collagen viscoelasticity |  |  |
| --- | --- | --- | --- | --- | --- | --- | --- |
| $E_a$ [J/mol] | $A$ [s<sup>-1</sup>] | $\beta_d$ [-] | $\delta_d$ [-] | $\gamma_d$ [-] | $\mu_d^f$ [MPa] | $\phi$ [-] | $\tau$ [s] |
| $3.57 \times 10^5$ | $1 \times 10^{53}$ | 2.6 | 0.15 | 0.72 | 0.0015 | 10 | 50 |

### Three-state model

Our second model takes an alternative approach to the above model. While it retains the first element of the above model’s second state (i.e., change in stress-free configuration of coiled collagen), it additionally introduces a third state (i.e., second denaturation state). Thus, we first consider an initial stage of denaturation which leads to aforementioned coiling of the collagen fibers, which is, again, driven by a rate *k_c_* of Arrhenius-type with parameters *A_c_* and *E_ac_*. However, additionally, we assume that a process of damage (or breakdown) occurs subsequently with a rate *k_b_*, also of Arrhenius-type with parameters *A_b_*, *E_ab_*. This breakdown leads to a reduction in collagen’s energy-storage capacity. Thus, as before, let *C*_o_ denote the intact collagen concentration. Additionally, we introduce *C*_c_ to denote the concentration of coiled collagen and *C*_b_ to capture the damaged collagen concentration. As a result, the strain energy of this mixture reads

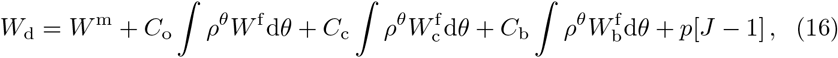

where 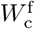 and 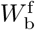 are the strain energy contributions due to the fibers after coiling and after being additionally damaged, analogous to Equations (4) and (11). Note that these energies are driven by their corresponding shear moduli 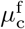 and 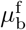, and waviness parameters *γ*_c_ and *γ*_b_, respectively. The respective second Piola-Kirchhoff stress tensors for the individual fibers and for skin follow identically to the previous derivations. All parameters for the three-state model – except for those that are identical to intact skin – are summarized in Table 3. As before, some of these parameters are estimated based on the literature, e.g. the rate of denaturation, while others are adjusted manually to approximate the results of Part I.

**Table 3:** Parameters for the three-state model of skin’s thermo-mechanical response to super-physiological temperatures

| Kinetics |  |  |  |  |
| --- | --- | --- | --- | --- |
| $E_{ac}$ [J/mol] | $A_c$ [s<sup>-1</sup>] | $E_{ab}$ [J/mol] | $A_b$ [s<sup>-1</sup>] | |
| $3.57 \times 10^5$ | $1 \times 10^{53}$ | $0.5 \times 10^5$ | $6.8 \times 10^5$ | |
| Waviness |  |  | Collagen Moduli |  |
| $\beta_c, \beta_b$ [-] | $\delta_c, \delta_b$ [-] | $\gamma_c, \gamma_b$ [-] | $\mu_c$ [MPa] | $\mu_b$ [MPa] |
| 2.6 | 0.15 | 0.72 | 0.18 | 0.0015 |

## Results

### Kinetics of denaturation

Starting with the two-state model, under the initial condition *C*_o_ = 1, *C*_d_ = 0, the analytical solution for the denaturation process reads simply as

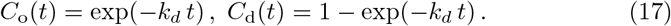

Solutions to above equation for temperatures *T* = 37, 55, 75, 95 °C are shown in Figure 2A-B. The model with our chosen parameters accurately predicts that denaturation is negligible at *T* = 37 °C. At *T* = 55 °C, in contrast, there is marginal decay in *C*_o_. As we increase the temperature to 75 °C drastic denaturation occurs with all collagen denaturing across the observation period of 100s. As we increase the temperature from 75 °C to 95 °C this process is nearly instantaneous. Figures 2C-D provide insight into the sensitivity of this process to the parameters of our model. Specifically, they show that increasing the activation energy *E_a_* significantly slows the denaturation process as measured by the time it takes for 50% of collagen to denature (ordinate). Conversely, increasing the rate parameter *A* significantly accelerates the denaturation process as measured, again, by the time it takes for 50% collagen to denature.

**Figure 2:**
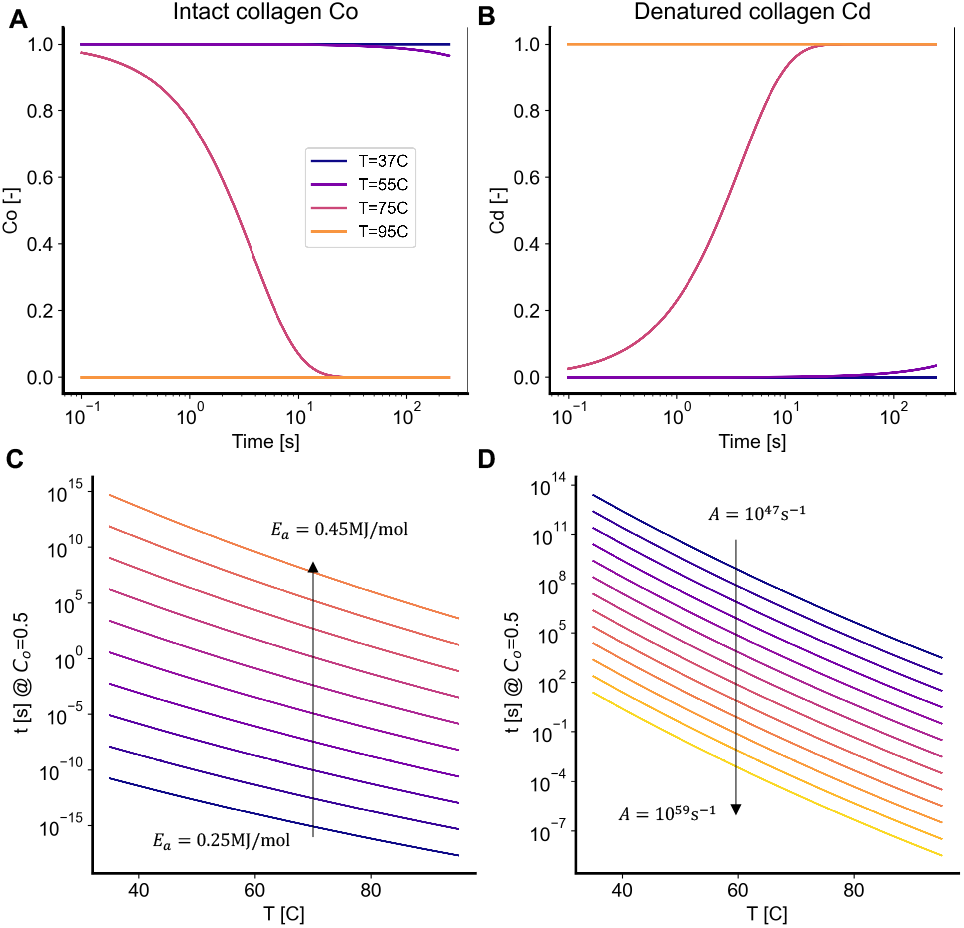
Denaturation kinetics of the two-state model. A,B) Concentration of intact (*C*_o_) and denatured (*C*_d_) collagen over time upon heat treatment at *T* = [37, 55, 75, 95] °C. C,D) Effect of activation energy *E_a_* and maximum rate *A* on the time it takes to reach half of the denaturation process (*C*_o_ = 0.5).

The kinetics of the three-state model are more complex, see Figure 3A. Here, the analytical solution for initial conditions *C*_o_ = 1, *C*_c_ = 0, and *C*_b_ = 0 reads

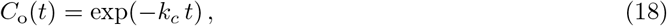

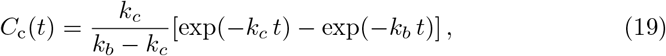

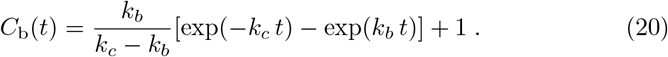

**Figure 3:**
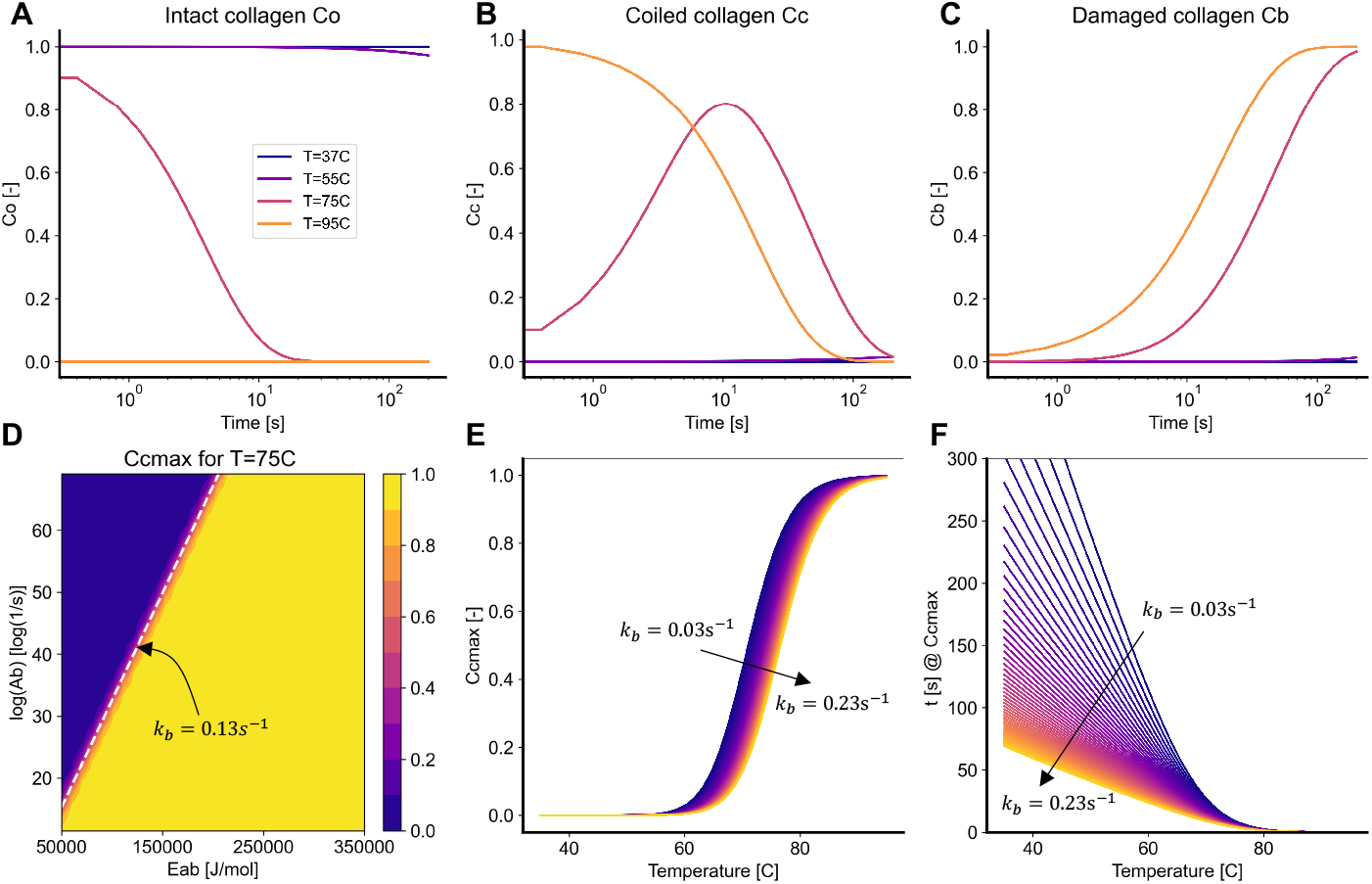
Denaturation kinetics of the three-state model. A) Concentration of intact (*C*_o_), B) coiled (*C*_c_), and C) damaged (*C*_b_) collagen over time upon heat treatment at *T* = [37, 55, 75, 95] °C. C) Contour of the maximum value that can be achieved by the coiled collagen concentration *C*_c*max*_ as a function of the secondary activation energy *E_ab_* and maximum rate of the secondary reaction *A_b_*. E) Effect of the rate of the second reaction (*k_b_*) on the peak of coiled collagen concentration *C*_c*max*_. F) Effect of the second reaction rate on the time to reach the peak *C*_c*max*_ as a function of temperature.

Note that the decay of the intact collagen follows the exact same time evolution as in the two-state model. However, in contrast to the two-state model, the three-state model of course tracks the temporal evolution of two concentrations of denaturing collagens; those of the second (coiled) state and those of the third (damaged) state, see Figure 3A-C. When heated to *T* = 75 °C, the coiled collagen concentration, *C*_c_, reaches a peak around 10s before decaying as *C*_b_ increases, i.e., collagen first coils and then damages. For *T* = 95 °C, *C*_c_ increases almost instantaneously and decays to zero over 100s. In other words, all collagen instantaneously coils upon which it quickly damages. Figure 3D-F clarifies the role of the parameters *A_b_*, *E_ab_* on the kinetics of the three-state model. An interesting feature of the three-state model is the maximum concentration of coiled collagen *C*_cmax_ because it, as we will see later, determines the peak in the thermally-induced traction production of skin, viz.

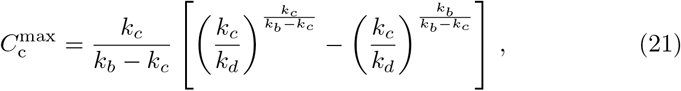

with the corresponding time at which the maximum coiled collagen is ob-served

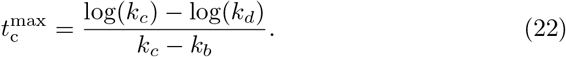

Figure 3D shows the contour of 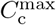 as a function of the parameters *A_b_*, *E_ab_* which control the second reaction in the three-state model. This contour shows that the peak of coiled collagen is very sensitive to the rate of the second reaction. For the majority of the parameters, the rate of the second reaction is either too slow – such that there is enough time for the native collagen to be converted almost entirely to the coiled state before transitioning to the final state – or too fast – such that there is no coiled collagen accumulation. There is only a narrow band in the *A_b_*, *E_ab_* over which the coiled collagen can take intermediate values. Setting *C*_cmax_ = 0.5 and solving equations (21) and (21), we obtain *k_b_* = 0.1302 s^−1^. The contour in Figure 3D then shows the line in the *A_b_*, *E_ab_* for *k_b_* = 0.1302*s*^−1^. The other two plots in Figure 3E-F then show how changes in *k_b_* influence the maximum value of *C*_c_ as well as the timing at which this peak occurs for a given temperature. Note that the curves in Figure 3E are similar in shape but shifted along the abscissa with a change in *k_b_*. For low temperatures there is no denaturation and no peak in the coiled collagen. As temperature increases, the rate of the first denaturation reaction increases which fuels an increase in *C_c_*, leading to the S-shaped curves in Figure 3E. However, accumulation of coiled collagen depends on *k_b_*, as *k_b_* increases there is less accumulation of *C_c_* and thus a lower peak for a given temperature. The time to reach the peak of coiled collagen decreases with an increase in *k_b_*. As illustrated in Figure 3F, the process is temperature-dependent. For high temperatures the peak occurs almost instantaneously independent of the second denaturation rate.

### Traction-free denaturation

In our first numerical experiments we directly mirror the experiments in Part I of this study by subjecting skin to four temperatures *T* = 37, 55, 75, 95 °C, in the absence of external tractions. Figure 4A shows the thermally-induced free shrinkage of skin according to the two-state model, while Figure 4B shows the results according to the three-state model. In both models, skin shrinks as the concentration of coiled collagen increases. Mathematically, the transition from native collagen to coiled collagen corresponds to a reduction in each fiber’s reference length, thus producing forces that, in the absence of external tractions, lead to shrinkage. Because of the fibers preferential distribution in lateral direction for mouse dorsal skin (i.e., 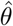 of the von Mises distribution equal to zero), forces are larger in that direction, thus leading to anisotropy in the free shrinkage. Following an initial contraction, the two-state model predicts a relaxation step due to the assumed viscoelasticity of the damaged tissue. The three-state model qualitatively predicts a very similar, overall behavior with negligible quantitative differences to the two-state model. Very interestingly, the three-state model also predicts a recovery of the initial shrinkage. However, rather than being due to viscoelastic recovery as in the two-state model, it is the transition from the coiled collagen to the damaged collagen that explains the loss in contraction. Since the properties of the damaged collagen *C*_b_ in the three-state model are the same as the equilibrium properties of the denatured collagen *C*_d_ in the two-state model, both frameworks lead to the same steady state response even though they follow somewhat different paths to equilibrium.

**Figure 4:**
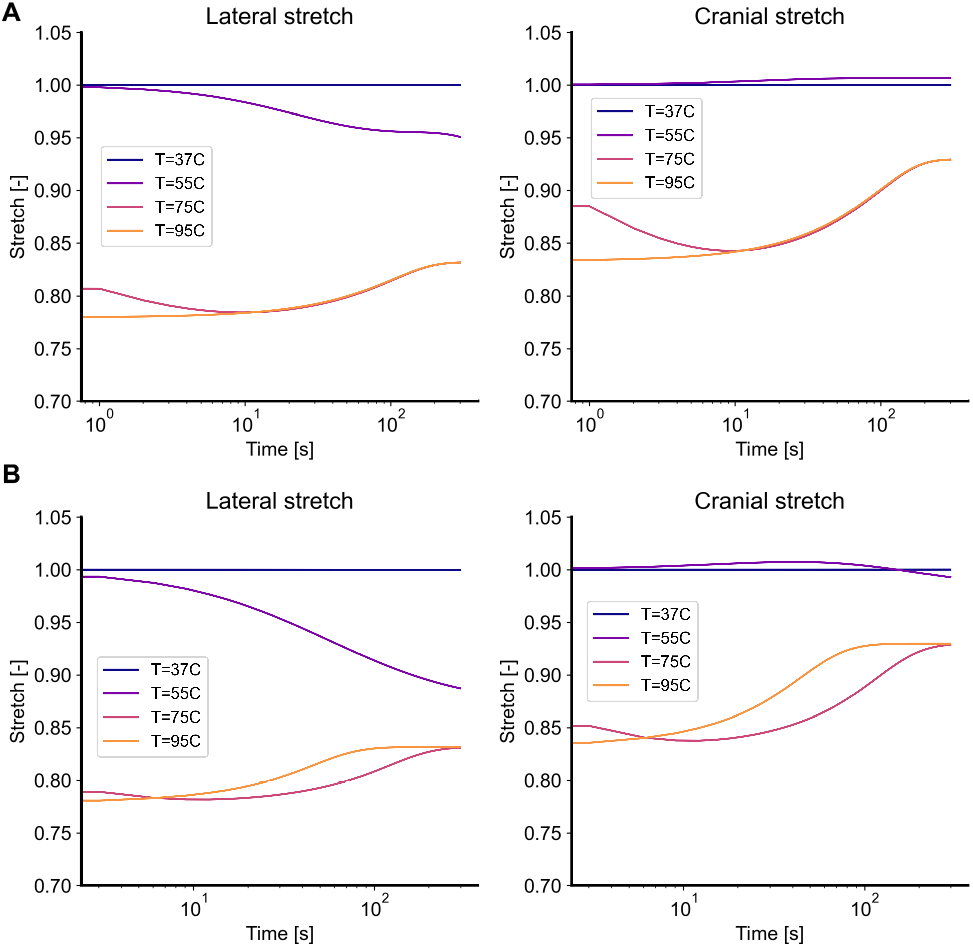
Traction-free shrinking for the (A) two-state model and (B) the three-state model. Upon subjecting it to temperatures of *T* = [37, 55, 75, 95] °C, skin shrinks over time. Both models predict that this free shrinkage is anisotropic, with greater deformation in the lateral direction than in the cranial direction. Additionally, both models predict that, after the initial shrinkage, skin also undergoes relaxation during which some of the shrinkage is recovered. In the two-state model this relaxation is due to the assumed viscoelastic behavior of denatured collagen, while in the three-state model this relaxation is due to the continuously accrued damage of the coiled collagen.

### Isometric denaturation

Again, in accordance with our experiments in Part I of this study, the next numerical experiment explores skin’s thermo-mechanics under isometric heating; that is, while skin’s boundaries are fixed. Both, the two-state model and the three-state model predict that isometric heating induces tractions that are higher in lateral direction than in cranial direction, see Figures 5A-B and 6A-B. Moreover, both models predict that those tractions are larger at 95 °C than at 75 °C and that – upon reaching a peak at approximately 10 seconds – those tractions decay and mostly vanish. However, note that the underlying physics are distinctly different. In the two-state model, forces are produced as the denaturing collagen is hindered from coiling, i.e., taking on a shorter reference length, while the subsequent decay follows from the viscoelastic relaxation of the denatured fibers. While the three-state model produces forces similarly as collagen transitions from the first state to the coiled, second state, the force decay mechanism differs. Specifically, in the three-state model the initial peak in force decays as collagen transitions from the coiled, second state, to a damaged, third state.

**Figure 5:**
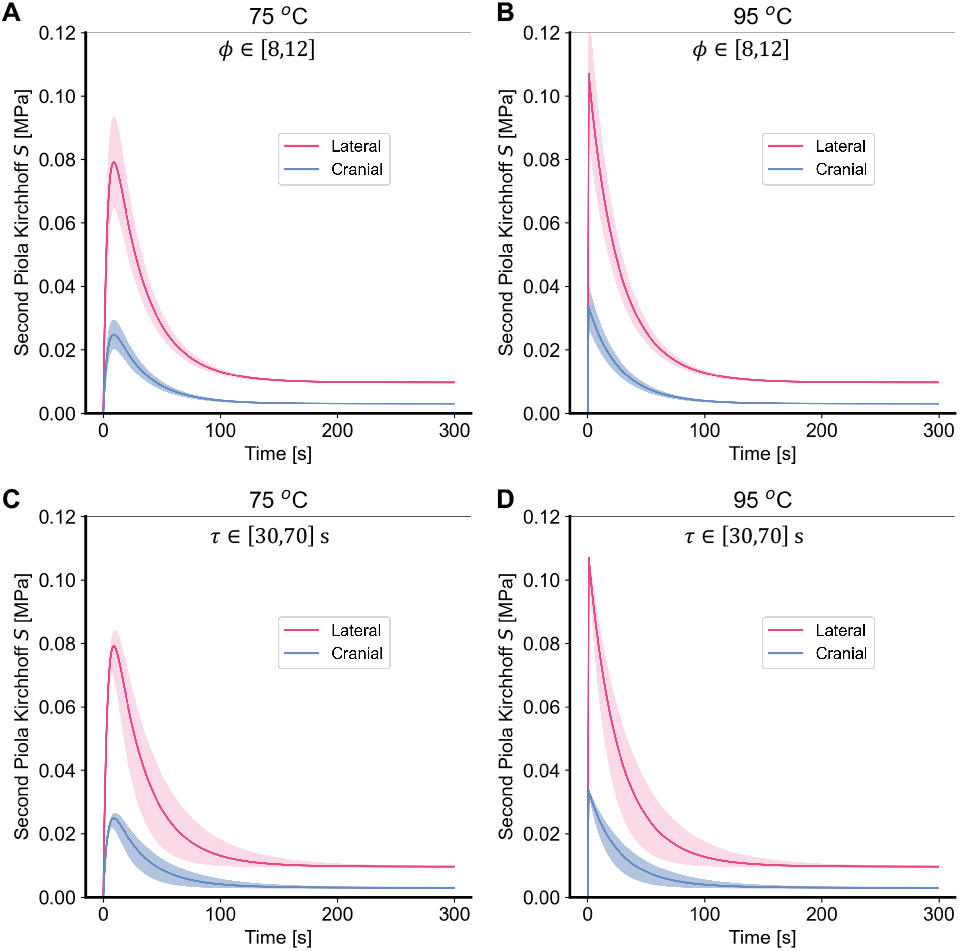
The two-state model predicts that skin produces tractions under isometric conditions and exposure to superphysiological temperatures. After an initial peak, those tractions decay and eventually mostly vanish. A-D) Effect of the parameter *ϕ* and *τ* of the two-state model at A,C) 75 °C and B,D) 95 °C, respectively, see Equation (12).

Figure 5C-D further explores the effect of critical model parameters of the two-state model: The parameter *ϕ* in Equation (12) controls the scaling of the dissipative branch in the viscoelastic model. Consequently, this parameter mostly influences the peak stress in Figure 5A,B. In contrast, the parameter *τ* in Equation (14) controls the timescale of stress relaxation. Thus, it primarily controls the decay time.

Figure 6C-D similarly explores the effect of critical model parameters of the three-state model. The peak and decay of the stress are primarily influenced by the modulus of the coiled collagen, *μ*_c_, and the reaction rate, *k_b_*, between the coiled, second state and the damaged, third state.

**Figure 6:**
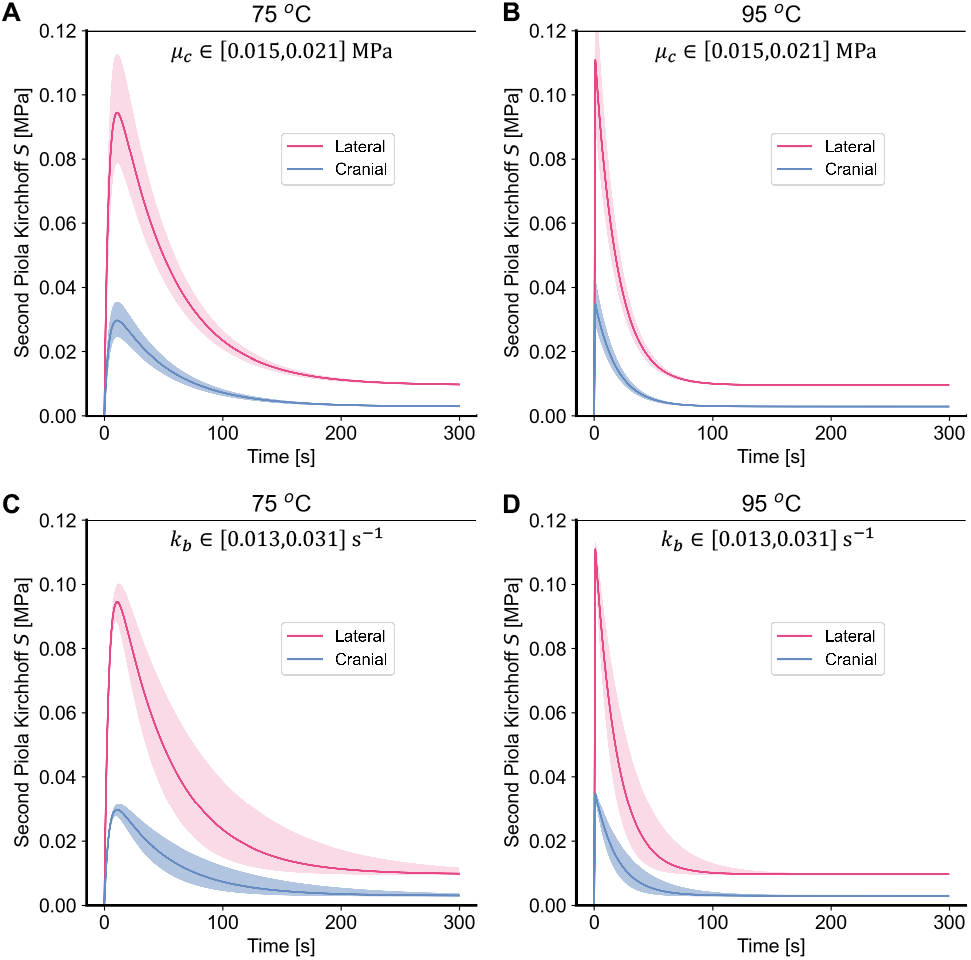
The three-state model predicts that skin produces tractions under isometric conditions and exposure to superphysiological temperatures. As with the two-state model, after an initial peak, those tractions decay and eventually mostly vanish A-D) Effect of the coiled collagen stiffness parameter *μ*_c_ and the reaction rate *k_b_* between the coiled collagen and damaged collagen of the three-state model at A,C) 75 °C and B,D) 95 °C, respectively.

### Biaxial mechanics following thermal denaturation

In the last numerical experiment we test skin’s constitutive response to 10s of thermal loading at 90 °C. Specifically, we compare skin’s biaxial mechanics before and after heat treatment. We thereby, once again, mimic our experimental protocol in Part I of this study. Both, the two-state model and the three-state model, predict that skin’s mechanics fundamentally change during denaturation, see Figure 7A,B for the results of both models, respectively. While skins biaxial response remains anisotropic, it changes from it’s classic J-shape – or strain-stiffening – behavior to a near-linear behavior. With this transformation, skin’s stiffness at small stretches increases, while it decreases at large stretches. The underlying, microstructural mechanisms are, in-fact, identical between the two models. That is, in both models the shift in the stress-free length of collagen molecules towards a shorter reference length shifts the “activation” or “uncrimping” length of collagen to smaller stretches; thus, increasing stiffness at small stretches. Simultaneously, coiled collagen is endowed with smaller shear moduli; thus, reducing stiffness at larger stretches.

**Figure 7:**
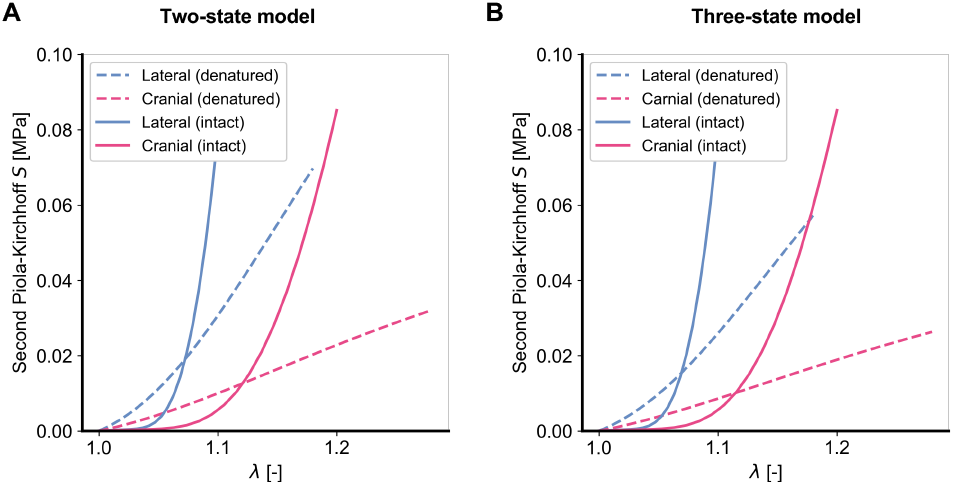
Biaxial properties of skin after 10s of denaturation at 90C, for the two-state model (A), and the three-state model (B). In both cases, denaturation leads to a stiffening at small strains but softening at larger strains.

## Discussion

In our current work we propose two competing models to capture the thermo-mechanical response of skin to superphysiological temperatures. This work was motivated by a lack of modeling and simulation tools that could explain our observations in Part I of this study that: i) skin shrinks anisotropically when heated under isotonic traction-free conditions, ii) skin generates anisotropic tractions when denaturing under isometric, or fixed-boundary, conditions, iii) and, when heated, skin changes its biaxial constitutive response to become more linear, with a stiffness increase at small stretches and a stiffness decrease at large stretches [21]. Our models build upon existing evidence that skin’s thermo-mechanical response to superphysiological loading is driven by collagen denaturation that follows Arrhenius-like kinetics [22]. Additionally, they are built on the knowledge that denatured collagen changes its stress-free reference configuration and that collagen changes its mechanical properties upon denaturation [20]. Our first, two-state model accounts for skin’s response through a transition from a first, native state to a second state in which collagen coils, i.e., takes a shorter stress-free configuration, and in which collagen transforms from a hyperelastic material to a hyper-viscoelastic material. In our three-state model, the second state also leads to coiling of collagen, i.e. reduced reference length, but does not lead to a transition to a hyper-viscoelastic material. Instead, in our three-state model, a third state leads to collagen damage through which its ability to store strain energy is compromised.

### Modeling success

In a first step toward modeling skin’s thermo-mechanics, we established a microstructurally-inspired hyperelastic model of skin’s native behavior. This baseline model builds on an explicit consideration of skin’s primary structural constituent: collagen. Specifically, we model collagen’s orientation and waviness according to probability density functions [32, 27]. This approach has successfully captured the anisotropy – via collagen’s orientation distribution – and the nonlinear stiffening behavior – via collagen’s waviness distributions– of other collageneous tissues such as arteries and thrombus [27, 33]. Among the advantages of this approach over other approaches, such as the projection of stress from the fiber network onto one or two structural tensors [26, 23], lies in our ability to bestow collagen fibers with microscopy-based measurements of orientation and waviness [29]. Our work shows that this modeling approach successfully captures all salient features of skin’s experimentally observed, bi-axial constitutive behavior. That is, our model correctly captures native skin’s anisotropy and strain-stiffening behavior.

Our two models of skin’s thermo-mechanical response to superphysiological loading share the same intact skin model. However, they differ by their accounting for a set of observations made in Part I of this study. Regardless of approach, numerical experiments with both models accurately reproduce all of our physical experiments. In the first numerical experiment, our models accurately predict that skin minimally shrinks under traction-free conditions when exposed to temperatures of 37 °C or 55 °C, but drastically shrinks when exposed to temperatures of 75 °C and 95 °C. Furthermore, they also correctly predict that the rate of shrinkage increases as a function of temperature. Interestingly, both models predict that, after an initial monotonic phase of shrinkage, skin expands again, a phenomenon that we did not identify in our experiments. The mechanisms underlying this late expansion differ between the two models. In our two-state model, the late expansion of skin is due to collagen’s transition to a hyper-viscoelastic material. In other words, after collagen coiling produces sufficient internal tractions to lead to shrinkage, these stresses then viscoelastically relax as strain energy dissipates. In our three-state model, the late expansion of skin is due to collagen’s transition from the second, coiled state to a third, damaged state of collagen. In other words, shrinkage follows from collagen coiling and build-up of internal tractions, while its late expansion results from a loss in collagen’s capacity to retain those internal tractions while it is becoming increasingly damaged.

In the second numerical experiment, our models accurately predict that skin produces tractions when its boundaries are fixed and it is subjected to 75 °C and 95 °C. Moreover, both models predict that those tractions rapidly decay after reaching a temperature-dependent peak. The traction production in both models stems from the transition of skin’s collagen from a native state to a coiled state. In other words, as collagen’s reference configuration becomes shorter, fixed boundaries lead to an induction of strain energy and thus tractions. However, both models differ in how they capture the subsequent decay. In the two-state model, the decay is due to viscoelastic relaxation of collagen. On the other hand, the three-state model captures the decay via collagen damage. In both models, the direction dependence of the traction peak stems from the relatively higher distribution of collagen in the lateral direction than in the cranial direction for mouse dorsal skin. Also in both models, the temperature-dependence of the traction peak is due to the temperature-accelerated coiling of collagen relative to the characteristic time of each model’s respective decay mechanism.

In the third numerical experiment, we compared skin’s constitutive response before and after being exposed to 10s of 90 °C. Here, again, both models accurately predict our experimentally observed phenomena. Specifically, both models accurately predict that skin transitions from a J-shaped, strain-stiffening constitutive behavior to a (mostly) linear constitutive behavior. That is, skin becomes stiffer at small stretches and less stiff at large stretches. In contrast to previous phenomena where the underlying mechanisms between the two-state model and the three-state model differed fundamentally, they don’t in this experiment. The reason being that the relatively short exposure to superphysiological loading of 10s leads to skin’s response in both models being governed primarily by collagen coiling. For example, in the two-state model the viscoelastic time constant *τ* is 50s. In other words, after 10s of exposure collagen’s behavior is still primarily hyperelastic. Similarly, the reaction rate of the three-state model that governs the transition from the second to the third state results in there being almost no collagen in the damaged state at 10s, see Figure 3. Consequently. both models predict the temperature-induced change in constitutive behavior based on collagen coiling and the associated reduction in collagen’s stress-free configuration and stiffness.

Please note that we described our modeling success primarily in qualitative terms. In fact, we did not conduct a formal fitting between our model and our experiments. The reason lies in our motivation to test our models’ basic, mechanistic-assumptions rather than to calibrate our models. Nonetheless, we manually chose our models’ parameter values so that our quantitative predictions match our experimental values closely.

### Limitations and future work

While both of our models are based on microstructural and molecular considerations, they don’t capture the molecular scale directly. Further numerical– and experimental – work on the molecular scale is needed to test our proposed mechanisms and to refine our predictions at the tissue level. Potential molecular-scale numerical methods that have previously been applied are coarse-grained methods, statistical mechanics, or molecular dynamics by Stylianopoulos et al., Miles et al., Yeo et al., and Mlyniec et al., [20, 34, 35, 36]. Such simulations may provide additional insight into the main mechanisms driving our models, such as collagen coiling, damage, and inter-fibril sliding. Thereby, they may shed light, at least theoretically, as to which model is more accurate and also inform future experiments. Beyond our lack of molecular insight, another significant limitation of our work is the reduction of the present thermo-mechanical problem to a homogeneous one. For example, we ignore heat transfer phenomena and assume instantaneous thermal equilibrium. In reality, the transfer of heat from the reservoir to the tissue may not be instantaneous, and there are likely gradients of temperature between the edge and the center of the tissue. We justify neglecting these spatial effects with our samples’ small dimensions of approximately 10mm 10mm 1mm. Nevertheless, additional work is necessary to test these assumptions – for example following the framework outlined by McBride et al. [37]. We also made simplifying assumptions about the constitution and constitutive behavior of skin. That is, we ignored native skin’s viscoelasticity/poroelasticity and heterogeneous composition [38, 39, 40, 41]. Finally, we did not consider more complex interactions such as a potential coupling between loading and thermal stability [42], or that the thermo-mechanical response may also be driven by other proteins, specifically elastin. Here, again, future work will have to explore these protein’s relative contribution to skin’s thermo-mechanical response to superphysiological loading.

## Conclusion

In our current work we propose and compare two models that link the kinetics of collagen denaturation to skin’s coupled thermo-mechanical response under superphysiological temperatures. Both models capture all experimental tissue-level observations that we made in Part I of this study. Because both models fundamentally differ in their underlying microstructural and molecular mechanisms, they may inform future experiments that could test each of the models’ assumptions. Additionally, our models may be used predictively. Therefore, our models are a crucial step toward a deeper understanding of thermal skin injuries and toward the rational, model-based design of skin’s – and other collageneous tissues’– thermal treatments.

## Author Contributions

William Meador designed and conducted all experiments, co-authored the manuscript. John Toaquiza Tubon developed the mathematical framework of native skin mechanics and co-authored the manuscript. Omar Moreno-Flores developed the mathematical framework of native skin mechanics and co-authored the manuscript. Adrian Buganza Tepole designed the experiments, developed the mathematical framework, conducted all simulations, and co-authored the manuscript. Manuel Rausch designed the experiments, developed the mathematical framework, and co-authored the manuscript.

## Acknowledgments

We acknowledge the National Science Foundation for their partial support of this project via Grant #1916663 (Rausch, Buganza).

## Supplementary Material

The code needed to reproduce all the figures in this paper can be found at https://github.com/abuganza/skin_thermomechanics_model.

